# The succession pattern of bacterial diversity in compost using pig manure mixed with wood chips analyzed by 16S rRNA gene analysis

**DOI:** 10.1101/674069

**Authors:** Zhengfeng Li, Yan Yang, Yuzhen Xia, Tao Wu, Jie Zhu, Zhaobao Wang, Jianming Yang

## Abstract

The pig manure mixed with wood chips and formed compost by means of fermentation. We found that the protease activity, organic matter content and ammonium nitrogen concentration were higher in the early stage of composting. Meanwhile, the urease activity was highest in the high temperature period. The carbon to nitrogen ratio of the compost decreased continuously with fermentation. The dynamic change in the composition of bacterial overtime in the compost of a 180 kg piles were explored using microbial diversity analysis. The results showed that the microbial species increased with the compost fermentation. At the early stage of composting, the phyla of Firmicutes and Actinomycetes were dominant. The microbes in the high temperature period were mainly composed of Firmicutes and Proteobacteria while the proportion of *Bacteroides* was increased during the cooling period. In the compost of maturity stage, the proportion of *Chloroflexi* increased, becoming dominant species with other microorganisms including *Firmicutes, Proteobacteria, Bacteroides, Chloroflexi* but not *Actinomycetes*. Bacteria involved in lignocellulose degradation, such as those of the *Thermobifida, Cellvibrio, Mycobacterium, Streptomyces* and *Rhodococcus*, were concentrated in the maturity stages of composting. Through correlation analysis, the environmental factors including organic matter, ammonium nitrogen and temperature were consistent with the succession of microbial including *Rhodocyclaceae, Anaerolineaceae, Thiopseudomonas, Sinibacillus* and *Tepidimicrobium*. The change of urease activity and carbon to nitrogen ratio corresponded to microbial communities, mainly containing *Anaerolineaceae, Rhodocyclaceae, Luteimoas, Bacillaceae, Corynebacterium, Bacillus, Anaerococcus, Lactobacillus, Ignatzschineria*, and *Bacillaceae*.

## Introduction

Aerobic composting of livestock manure and agricultural waste is the most economical and environmentally friendly way of obtaining a fertilizer. During this process, most microbes grow under aerobic conditions. Compared with anaerobic fermentation, the aerobic fermentation cycle is shorter. The resulting composts can be used in farmland as biological organic fertilizers, which is of great significance for promoting ecological agriculture.

Aerobic composting generally undergoes four stages, including the heating period, high temperature period, cooling period and maturity period Although the microorganism communities of compost are highly complex, the succession of microbial communities during composting obeys certain rules [1]. The traditional research methods analyzing the microorganisms composition in compost mainly include PLFA, DGGE, PCR-RFLP and plate culture method, etc.[2] However, because of the limitations of culture conditions and the low resolution of electrophoresis gels, these analyses of microorganisms have not been comprehensive. Currently, the 16S RNA special fragment is amplified using the bacterial and archaeal primers. By comparing amplification fragment of 16S RNA with online databases of bacteria sequence, the microorganism’s type and quantity in the compost are determined [3].

Traditional composting methods include composting with single raw materials and mixed raw materials. The fundamental reason for using various additives in composting processes is to provide the best-growing environment for microbial growth and achieve rapid composting, thus reducing the harmful gas emissions under controlled conditions. For example, sawdust is an additive that reduces methane emissions to a certain extent [4]. However, the quality of the fertilizer ultimately depends on the microbial composition of the compost, and only a clear understanding of the succession pattern of the microbial communities throughout the composting stages will enable us to fully control this process. This knowledge would help us find the best compost additives and microbial agents, laying the foundations for screening microorganisms for special functions.

In this study, we studied the evolution of bacterial communities in a mixed compost of pig manure and sawdust under the condition of a low carbon to nitrogen ratio (20-25:1). Using 515F and 806R primers, the 16S RNA specific region of prokaryotic genomes was amplified, and the whole process of composting bacteria was systematically investigated. By analyzing the changes in compost of the dominant bacteria and the correlation between microbial community and important environmental parameters, we managed to explain the effects of environmental factors on the changing microbial communities in compost and provide scientific advice to control the fermentation process of the compost by adjusting the physicochemical parameters.

## Materials and methods

### Compost composting and sampling

The compost study was performed in three piles (diameter, 1.5 m; height, 1.1 m). An 800 cm length, 20 cm wide, 20 cm deep trench was dug at the surface and three 200 cm length, 20 cm wide and 20 cm deep trenches were used to vent at the bottom of the compost. The branches and wheat straw were alternately put in the cross position of each groove. Then the compost with pig manure and sawdust mixed with C/N ratios of 25:1 was laid on top. A stick was inserted at the top of a pile of compost to increase the overall aeration, and the piles were covered with a black perforated plastic sheet to avoid heat and water loss. Three thermometers were inserted in different directions into the 75 cm high compost. The real-time temperature of each compost stack was the average value from three different sites. Water was added during the experiment to maintain a moisture content level of 60 %. The oxygen was supplied during the composting process by turning the pile once every five days.

Through long-term composting test, the 50 days of the natural fermentation process samples in different fermentation stages were studied. Firstly, we determined the midpoint of the diagonal as the central sampling point. Secondly, we selected four additional samples equidistant from the center point. These five sub-samples were then fully mixed to form one composting sample. Three samples from three piles were obtained at each fermentation stage, and 12 samples were obtained at four fermentation stages including the prime stage, high-temperature period, cooling period, and maturity period. Samples obtained from each composting were labeled Z31, Z32 and Z33 for third days after composting. The compost samples taken on day seven were referred to as Z71, Z72 and Z73. The samples taken at day 20 were labeled Z201, Z202 and Z203. On the fiftieth day, samples obtained from the composts were labeled Z501, Z502 and Z503. All samples were stored at −80 °C until use.

### Compost physical and chemical analysis

The temperature was measured every day in the compost by thermometer place at the same height and depth. Measurements were taken from three thermometers placed at different angles. The water extract with a 1:4 ratio of the compost sample contrast to distilled water was used for the measurement of pH. The final pH was the average of three measurements. Total ammonia nitrogen was analyzed by the MgO distillation method [5]. The total organic carbon was determined according to Walkley-Black wet combustion method [6]. The total organic matter was the number of total organic carbon multiplied by 1.725. The total N was determined by the previous Kjeldahl method.

### Protease activity and urease activity assays

Protease enzyme activity was assayed by Casein Digestion Method [7]. For determination of urease activity, the method of magnesium oxide distillation was used [8].

### DNA extraction and high throughput sequencing

DNA was extracted from 0.5g of compost sample (wet weight) using a FastDNA^®^ Spin Kit for Soil (MP Biomedicals,USA), based on the manufacturer’s instructions. Primer sets 515F (5’-GTGCCA GCMGCCGC GG-3’) and 806R (5’-GGACTACHVGGGTWTCTAAT-3’) were used for bacterial 16S rRNA specific region amplification. During synthesis, barcode and adaptor sequences were added to the sequencing primers. An aliquot of a 20 μL PCR reaction included 4 μL 5× FastPfu buffer, 2 μL 2.5 mM deoxynucleoside triphosphate (dNTP) mix, 0.4 μL of each primer, 0.4 μL TransStart Fastpfu DNA Polymerase (TransGen), 1 ng template DNA and 11 μL double-distilled H_2_O. The PCR reactions were performed in triplicate under the following conditions: initial denaturation at 95 °C for 2 min, 25 cycles of 95 °C for 30 s, 55 °C for 30 s, 72 °C for 30 s, and then a final extension at 72 °C for 5 min. Each sample was amplified in triplicate, pooled and purified using AxyPrep DNA Gel Extraction Kit (AXYGEN). The concentration of purified PCR products was measured with QuantiFluor™-ST system (Promega). Then the barcode-tagged amplicons from different samples were mixed in equimolar concentration and sent to the Majorbio Bio-Pharm Technology Co., Ltd. (Shanghai, China) for Miseq library construction and sequencing. The raw sequence data were deposited into the NCBI short reads archive database (accession number: SUB3155541).

### Data preprocessing and bioinformatics analysis

The original fastq files have undergone quality filtering and data merging to prevent mismatch. Reads containing ambiguous bases were removed. Operational taxonomic units (OTUs) were clustered with 97 % similarity cutoff with a novel algorithm that performs chimera filtering and OTU clustering simultaneously. These sequences are bound to the operation classification unit (OTUs) with 97 % recognition television threshold. The most abundant sequence in each OTU was selected as the representative sequence. The 16S rRNA gene was screened by RDP Classifier algorithm against the Silva (SSU123) 16S rRNA database with a confidence threshold of 0.7.

### Statistical analysis

Based on the results of OTU cluster analysis and taxonomic information, a series of in-depth statistical and visual analysis on community structure and phylogeny such as OTU generation, sampling adequacy analysis, abundance and diversity analysis, flora difference analysis, evolutionary tree analysis, etc. could be carried out to screen samples with larger microbial abundance or more complex community structure.

For Illumina Miseq sequencing data, alpha diversity indices, Chao, ace, Shannon index and inverse Simpson index were calculated using the Quantitative Insights Into Microbial Ecology (QIIME). In the beta diversity analysis, the weighted UniFrac distance and Bray-Curtis distance were calculated using the “pure prime” packets by QIIME and ‘vegan’ package in ‘R’, respectively. Principal coordinates analysis was conducted to visualize the community similarity with the ‘vegan’ package in ‘R’. Linear Discriminant Effect Size (LEfSe) analysis was used to identify the microbial taxa that were significantly associated with each sample. The alpha value of Kruskal-Wallis test is 0.05, and the threshold value of Logarithmic Linear Discriminant Analysis (LDA) score is 2.0. Using Welch’s t test with Bonferroni correction in’ STAMP’, the differences of relative abundance among different samples were analyzed. The microbial community diversity indices, including number of sequences, Shannon-Wiener index and evenness index, were calculated as described before. Data were analyzed by analysis of variance (ANOVA). Alpha diversity index of Illumina Miseq sequencing was tested by Welch’s t test, and the mean value between samples was compared at a probability level of 0.05.

## Results and discussion

### Summary of 16S rRNA gene sequencing

In this study, the Illumina Hiseq 2500 platform was used to study the succession pattern of bacteria in different fermentation stages of compost. The bacteria diversity and quantity in samples from different fermentation stages are shown in Table 1. The order of the Shannon indexes of twelve samples was as follows, Z50 > Z7 > Z20 > Z3, which indicated that the bacteria diversities at the thermophilic phase and maturity stage were higher than that at the primary stage and cooling stage (Fig 1). However, the Chao1 index increased with the time of compost fermentation, which suggested that the quantity of microorganism was raised with compost fermentation.

**Fig 1.**
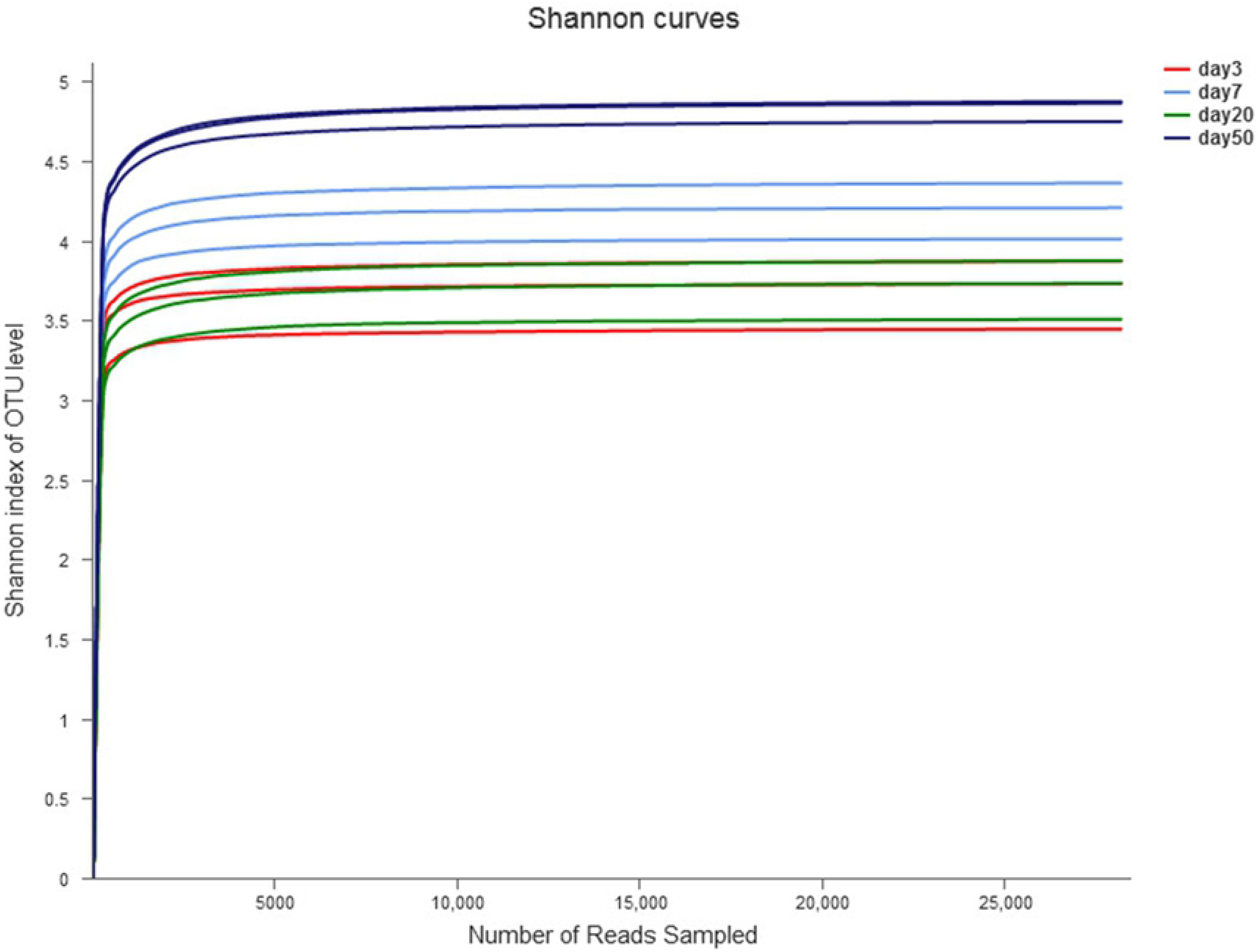
Shannon curves of samples with different days of fermentation.

**Table 1.** Sequences and sobs of the different fermentation stage of compost with 97 % similarity.

| sample | sequence | Shannon | Chao1 | coverage | OTU |
| --- | --- | --- | --- | --- | --- |
| Z31 | 34833 | 3.878 | 687.288 | 0.996 | 554 |
| Z32 | 40998 | 3.734 | 660.041 | 0.996 | 517 |
| Z33 | 34809 | 3.449 | 483.333 | 0.997 | 428 |
| Z71 | 44462 | 4.361 | 914.642 | 0.996 | 795 |
| Z72 | 35256 | 4.023 | 723.463 | 0.995 | 589 |
| Z73 | 32556 | 4.209 | 780.184 | 0.995 | 643 |
| Z201 | 35324 | 3.873 | 1010.505 | 0.994 | 813 |
| Z202 | 33229 | 3.738 | 978.632 | 0.994 | 797 |
| Z203 | 40320 | 3.514 | 848.072 | 0.996 | 689 |
| Z501 | 43669 | 4.865 | 1254.128 | 0.996 | 1133 |
| Z502 | 39968 | 4.879 | 1220.719 | 0.995 | 1066 |
| Z503 | 30433 | 4.749 | 1130.203 | 0.992 | 942 |

The relative abundance curves showed the evolution of microbial diversity in compost samples (as shown in Fig 2). The diversity of composting samples was the lowest at day three and the highest at day 50, indicating that high diversity was the stable status of the compost. The diversity of compost samples varied between days seven and twenty. As shown in Table 1, the richness was higher at day twenty than day seven. The diversity of microorganisms is affected by richness and evenness. Hence, the evenness of the samples from day seven was higher than that of the samples from day 20. This suggested that the seventh-day compost samples could be enriched for high-temperature resistant bacteria. Since the organic matter content was rich and other environmental factors had little influence on the microorganisms at the thermophilic phase, thermophilic bacteria capable of degrading organic matter grew heavily in the compost.

**Fig 2.**
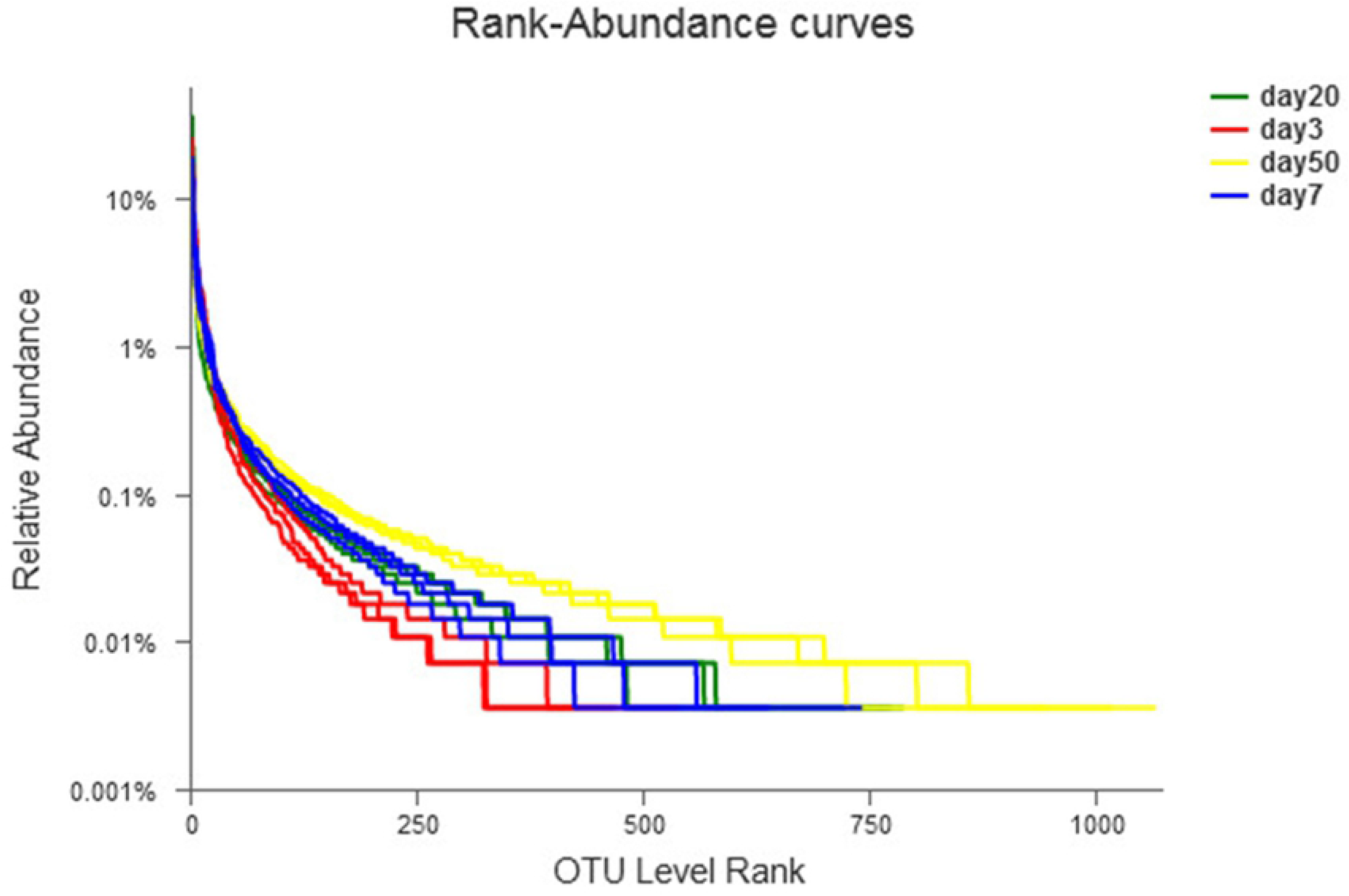
Rank-Abumdance curves of samples with different days of fermentation.

### Bacterial community changes at the phylum and genus levels

Seven phyla were detected in all samples. Of these, five phyla accounted for the vast majority of sequences and were considered dominant phyla: *Firmicutes*, *Proteobacteria*, *Bacteroides, Actinobactrria* and *Chloroflexi* (Fig 3). As the composting proceeded, a succession of genera was observed. From the primary stage (Day 3) to the late stage (Day 50), the abundance of *Firmicutes* decreased from 70 % to 26 %, while *Proteobacteria* and *Bacteroides* increased from 9.6 % to 28 % and 1.4 % to 22 %, respectively. *Firmicutes* could grow at high temperatures and was present in the thermophilic phase during composting of agricultural and sideline products [9], taking dominance at the early stages of composting. *Proteobacteria* and *Bacteroides*’ dominance was likely to exist at lower temperature in the maturity period. In the samples from the maturity period, the proportion of *Chloroflexi* was higher than in the other periods. Similarly, these dominant phyla also abounded in other lignocellulosic substrates compost [10].

**Fig 3.**
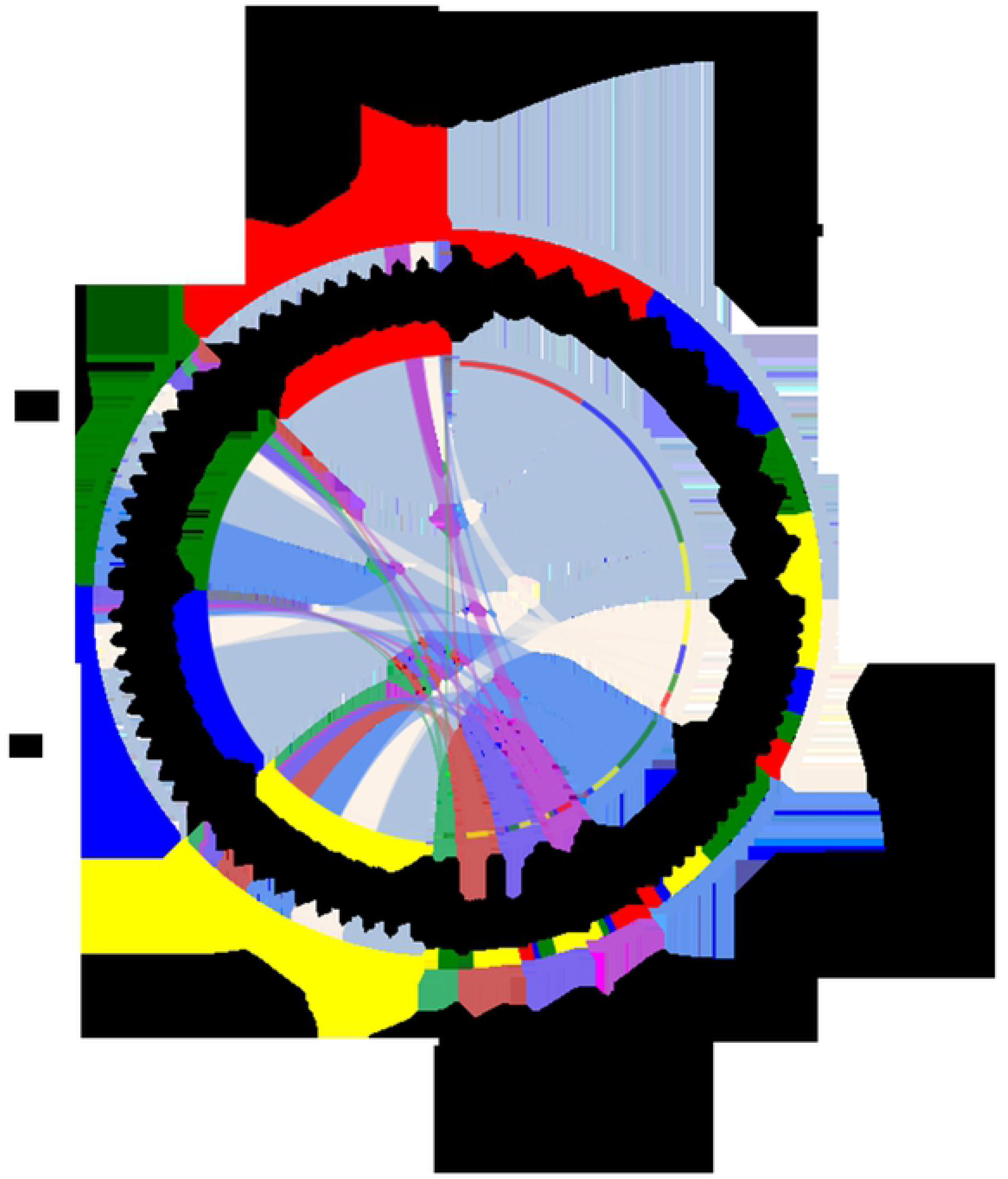
Distribution of microorganisms at bacterial phylum level in composting samples of different fermentation stages.

As showed in Fig 4, the change in the bacterial community diversity at the genus level was similar to that of the phyla level. At the initial stage of composting, genus including *Corynebacterium, Ignatzschineria*, *Lactobacillus, Sinibacillus* and *Thiopseudomonas* dominated in the community, but these were eventually taken over by genus such as *Ruminofilibacter*, *Hydrogenispora, Anaerolineaceae, Bacteria* and *Petrimonas* in the maturity stage. Microbial diversity at the genus level in the maturity stage was higher than that in the other fermentation periods. As the fermentation proceeded, the compost tended to stabilize in the diversity of genera.

**Fig 4.**
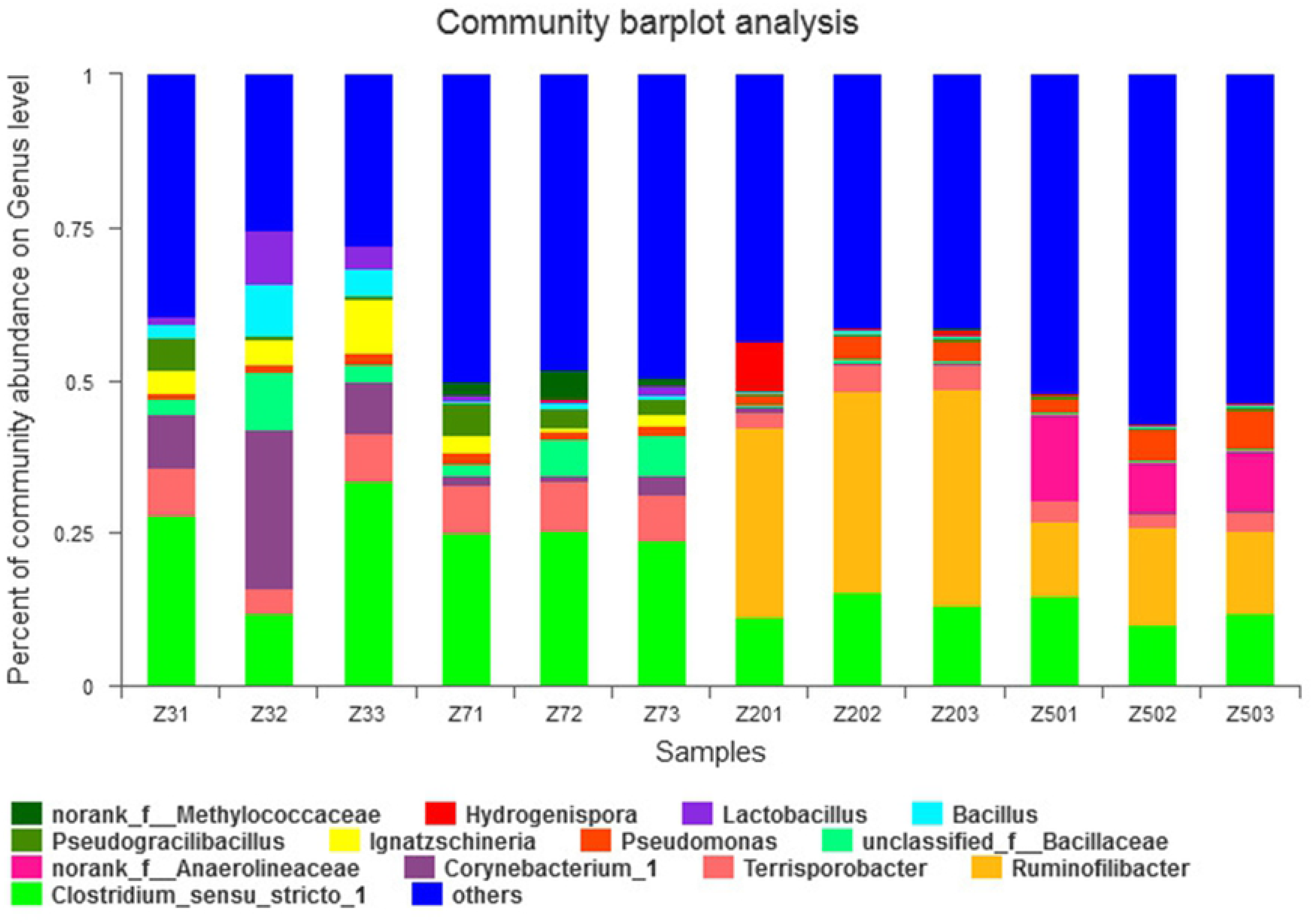
Genus level composition of the bacterial communities in the compost at different fermentation days (Merge bacteria less than 2 % into the others).

Sequences were considered to belong to different species if the sequence differences are higher than 2 %. The microorganisms in the early stage of composting decreased in the later stage were *Orynebacterium, Ignatzschineria, Lactobacillus, Bacillus, Sinibacillus* and *Thiopeudomonas*. As the compost went on, the microbes whose numbers had increased were mainly *Ruminofilibacter*, *Hydrogenispora* and *Petrimonas*. The changes in bacterial community composition during composting were illustrated using heatmap as in Fig 5. *Clostridium-sensu-stricto*, *Terrisporobacter*, *Turicibacter* and *Pseudomonas* did not change significantly during the composting process. The quantity of *Caldicoprobacter* increased by day seven and then decreased in the maturity period. Being in a thermophilic anaerobe group, its optimum growth temperature was 65 °C and the optimum growth pH was 6.9. The bacteria could withstand a high concentration of salt [11]. *Corynebacterium-1* only existed in the primary composting and high-temperature period, not in the cooling period and maturity stage. Some bacteria of this genus were pathogenic and were reduced in the late composting stage, indicating that composting played an important role in killing pathogenic microorganisms. The *Ignatzschineria* grew in an aerobic and low-temperature environment and then decreased gradually. As they had urease and phenylalanine deaminase activity [12], these bacteria mainly lived in the initial stage of composting that was an organic-rich and low-temperature environment. *Lactobacillus* was not resistant to high-temperature and high-concentration of salt and they could only grow in an anaerobic environment [13]. This was mainly due to the rapid degradation of organic matter in the early stage of composting, resulting in a partial anaerobic environment. As the temperature increased gradually, the easily degradable organic matter became scarce, particularly in late composting. *Bacillus* had strong amylase activity and could grow in various carbon sources [14]. With the ability to generate spores, *Bacillus* could sustain the high osmotic pressure and temperature in the compost. In the late composting stage, *Bacillus* decreased due to the reduction of organic matter in the compost. *Thiopeudomonas* was suitable for growing in an anaerobic and alkaline environment. It had the ability to hydrolyze lipid and denitrify the nitrate nitrogen. At the same time, It could also resist to high temperature [15]. Thus, its quantity was higher in the alkaline environment, with high ammonium nitrogen concentration at the early stage of composting. *Ruminofilibacter* could hardly be detected at the early stage of composting, but this group reached high numbers during the cooling period. This probably occurred due to its ability to degrade xylan [16–17], In the late composting, the main organic compounds were lignocellulose, which could be degraded into the xylan that sustained the growth of *Ruminofilibacter. Petrimonas* was intolerant to high temperature [18]. Accordingly, they increased during the cooling stage of the compost. *Clostridium spp*. were abundant in the whole composting process, accounting for a relatively large proportion in the early stage of composting, whereas their number decreased in the late stage of composting. These were anaerobic microorganisms that played an important role in the degradation of organic matter, including lipids, proteins and polymeric carbohydrates [19–20]. *Terrisporobacter*, which produced spores, could adapt to a high osmotic pressure environment [21]. Hence, this group was present during the whole compost process. *Turicibacter* was strictly anaerobic and remained abundant in the compost, indicating a lack of oxygen in the natural compost. *Pseudomonas*, an important lipid-degrading bacteria that dissolved minerals and provided nutrients [22], were highly productive at different stages of the compost. Therefore, these bacteria contribute to the quality of the compost as a fertilizer. Hill et al. found that *Bacteroides* had a very strong metabolic capacity for nutrients, such as complex organic matter, proteins and lipids [23]. These species were mainly concentrated in the early stage of the compost, and contributed to accelerating the decomposition of organic matter.

**Fig 5.**
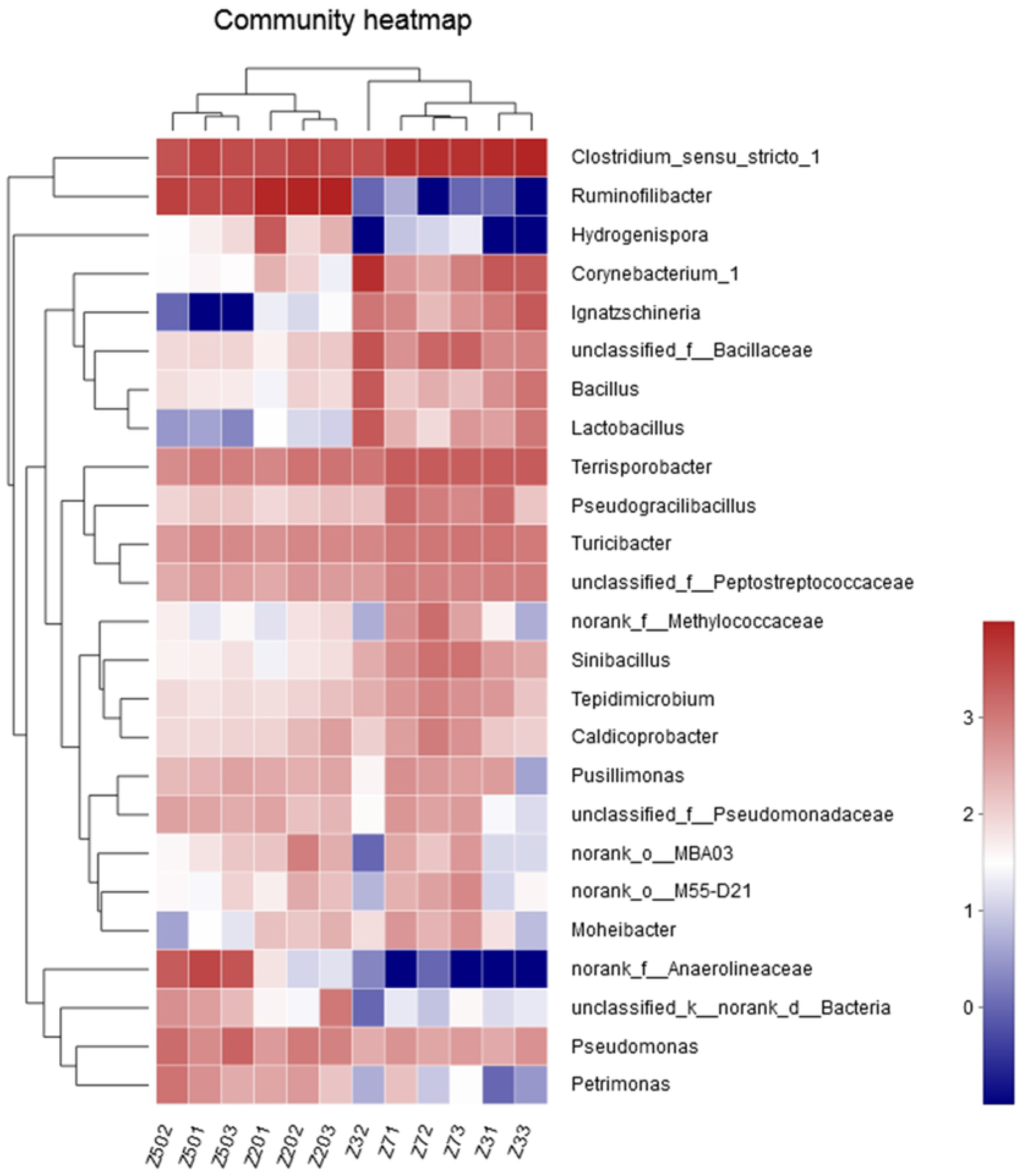
Heatmap illustrating the changes in bacterial community composition during composting. Each column is colored so that taxa with low abundances are green and high abundances are red.

### Lignocellulose-degrading bacteria associated with compost

Sawdust as a bulking agent was added to supply the extra carbon source, increasing the ratio of carbon to nitrogen. Specific microorganisms capable of lignocellulose degradation dominated the different stages of composting [9]. There are many lignin, cellulose and semi-cellulose components in sawdust, which could enrich the cellulose and lignin degrading bacteria. These included *Thermobifida fusca*, which in the matrix of lignocellulose compost could produce cellulase and hemicellulose degradation enzymes including xylanase, endo-1,4-beta-xylanase, beta-1,4- endoxylanase, endo-1,4-beta-xulanase and mannan endo-1,4-beta-mannosidae [24]. Consequently, it had the ability to degrade lignocellulose. As *Thermobifida fusca* tolerated high temperature, in combination with *Aspergillus*, had also been found to degrade lignocellulose [25], it was of great importance to degrade lignocellulose during the high-temperature period of composting. This species was detected during all stages of the composting process. *Cellvibrio*, a cellulose-degrading bacteria with low tolerance to high temperature, were found to be more abundant in the late stage of compost than in other periods [24]. Their presence greatly promotes the continuous degradation of cellulose.

Generally, in lignocellulosic compost, the main bacteria found were *Actinomyces*, including the genus of *Thermomonospora curvata*, *Mycobacterium xenopi*, *Amycolicicoccus subflavus*, and *Mycobacterium thermoresistibile*. Other bacteria that had been shown to degrade lignocellulose were *Streptomyces*, *Rhodococcus* and *Nocardia* [26]. In this study, the lignin-degrading bacteria of *Mycobacterium, Streptomyces*, and *Rhodococcus* genera were found, mainly in the late stage of composting and relatively less in primary phase. In conclusion, the microbes referring to the degradation of lignocellulose were enriched at the mature period. Because under the natural composting conditions, easily decomposed organic matters were abundant at the early stage of composting, which was not beneficial to the enrichment of these kinds of microbes.

*Cytophagaceae* was a family with the capacity of cellulose degradation function [27], which did not appear at day three of compost. At day seven of compost fermentation, *Pericitalea* was the dominant genus of *Cytophagaceae*. At day 50 of composting, the main genera of the *Cytophagaceae* family were *Chryseolinea*, *Ohtaekwangia* and some other genera without classification, showing different community structure. This indicated that lignocellulose was still degraded to some extent at the end of composting mainly by *Chryseolinea*, *Ohtaekwangia* some other genus.

During the composting process, the community structure of microorganisms showed significant succession. Temperature and nutrition played an important role in the microbial community structure succession. The bacteria developed from the primary superiority of microbial community capable of metabolizing simple organic matter to a mature microbial community characterized by the ability to produce lower amounts of biodegradable compound [28].

### Bacterial communities related to nitrogen and phosphorus metabolism

Nitrosation bacterium had the capacity for anaerobic denitrification. *Nitrosomonas* was more abundant at the end of composting [29]. Some other bacteria related ammonia oxidation to nitrite such as *Nitrosocystis*, *Nitrococcus* and *Nitrosospira* in compost were not detected. However, the amount of ammonium nitrogen decreased in the late stage. The reduction of ammonium nitrogen in the compost was mainly through the loss of volatilization. The whole composting process was not detected in nitrifying microorganisms such as *Nitrobacter, Nitrospina and Nitrococcus*. As chemoautotrophic bacteria, their frequency in the environment was relatively low and was very sensitive to the requirements of temperature, substrate and oxygen [30]. Judging from the quantity of other anaerobic bacteria in composting, most of the environment in composting was still anaerobic, which was not conducive to the growth of nitrifying bacteria.

The temperature had an important influence on the community structure of ammonification bacteria [31]. The population of ammonification bacteria in compost was numerous. The influence of external disturbance was very small. Although some species were decreased, at the same time other bacteria activity would rise to keep the balance, making the overall function of the microbial population relatively stable [32].

Genes related to the ammonization process comprise the main extracellular enzyme genes and intracellular enzyme genes, including alkaline metallopeptidase gene *apr*, serine peptidase gene *sub* [33]. Using the method of Southern-blot probe hybridization, Bach *et al*. revealed that *sub* gene was mainly found in a variety of *Bacillus spp*., and *apr* gene was mainly found in *Pseudomonas fluorescens*. The bacteria of these two genera could accelerate protein decomposition in the compost.

*Alcaligenes sp*. and *Sphingomonas sp*. were microbial species capable of ammonia removal [34]. *Sphingomonas sp*. was detected in the cooling period and the maturity period of composting, whereas *Alcaligenes sp*. was detected in the early period and the high temperature composting period. They had a synergistic effect of ammonium nitrogen metabolism, reducing the ammonium nitrogen loss during composting [34]. Some of the *Bacillus* genus had the ability of ammonia assimilation [6], previous studies also prove that *Bacillus smithii* had stronger ability of ammonia assimilation in nonsterilized compost extract media [35]. In this study, the *Bacillus* genus showed great abundance along the whole composting process as figure 5 described.

As the composting proceeds, the phosphorus in the compost was enriched. Thus, in the later stages of compost, the number of microorganisms related to phosphorus metabolism increased. *Anaerolineaceae* bacteria, belonging to *Chloroflexi* phylum, was a family of bacteria found in compost that related to phosphorus removal [36]. This group was found mainly at the end of composting, in agreement with its metabolic properties and the fact that phosphorus concentrations were high at the end of process.

At present, the most frequently reported phosphorus accumulating microorganisms (PAOs) in the genus were *Acinetobacter, Aeromonas, Corynebacterium*, and *Enterococcus* [37]. These bacteria existed in compost, having the effect on phosphorus enrichment. Notably, *Clostridium spp*. had denitrifying properties and the ability to remove phosphorus [38]. *Clostridium-sensu-stricto* was abundant throughout the composting process, which may be related to the phosphorus removal in whole compost process. *Anaerolineaceae*, which belonged to *Chloroflexi* phylum, could remove phosphorus [36] were found in the end stages of the process.

### Bacterial communities associated with environmental factors analysis

The relationships between chemical parameters and genus abundance (the dominant species of the top 35 in at least one sample) were evaluated in this study (Fig 6). The results showed that the urease (UE) correlated positively with the abundance of *Corynebacterium* (r = 0.893; P ≤ 0.001), *Bacillus* (r = 0.86; P ≤ 0.001), *Lactobacillus* (r = 0.916; P ≤ 0.001), *Ignatzschineria* (r = 0.96; P ≤ 0.001), *Anaerococcus* (r = 0.956; P ≤ 0.001), and negatively with the abundance of *Anaeralineaceae* (r = −0.861; P ≤ 0.001), *Rhodocycladeae* (r = −0.907; P ≤ 0.001), *Luteimonas* (r = −0.916; P ≤ 0.001). The C/N correlated positively with the abundance of *Bacillaceae* (r = 0.86; P ≤ 0.001), *Corynebacterium-1* (r = 0.925; P ≤ 0.001), *Bacillus* (r = 0.923; P ≤ 0.001), *Lactobacillus* (r = 0.944; P ≤ 0.001), *Ignatzschineria* (r = 0.949; P ≤ 0.001), *Anaerococcus* (r = 0.925; P ≤ 0.001), and negatively with the abundance of *Anaeralineaceae* (r = −0.854; P ≤ 0.001), *Rhodocycladeae* (r = −0.885; P ≤ 0.001), *Luteimonas* (r = −0.895; P ≤ 0.001). The apparent temperature [33] correlated positively with the abundance of *Bacillaceae* (r = 0.771; P ≤ 0.001), *Corynebacterium-1* (r = 0.716; P = 0.009), *Lactobacillus* (r = 0.782; P = 0.003), *Ignatzschineria* (r = 0.757; P = 0.004), *Anaerococcus* (r = 0.735; P = 0.006), *Caldicoprobacter* (r = 0.803; P = 0.002), *Thipseudomonas* (r = 0.917; P ≤ 0.001), *Sinibacillus* (r = 0.831; P = 0.001), *Tepidinicroblum* (r = 0.845; P = 0.001), and negatively with the abundance of *Rhodocycladeae* (r = −0.913; P ≤ 0.01), *Pseudomonas* (r = −0.754; P = 0.005). The concentration of NH_4_^+^ correlated positively with the abundance of *Bacillaceae* (r = 0.797; P = 0.002), *Corynebacterium-1* (r = 0.718; P = 0.009), *Lactobacillus* (r = 0.755; P = 0.005), *Ignatzschineria* (r = 0.722; P = 0.008), *Anaerococcus* (r = 0.732; P = 0.007), *Caldicoprobacter* (r = 0.818; P = 0.001), *Thiopseudomonas* (r = 0.888; P ≤ 0.01), *Sinibacillus* (r = 0.851; P ≤ 0.01), *Tepidinicroblum* (rr = 0.86; P ≤ 0.01), *Terrisporobacter* (r = 0.727; P = 0.007), and negatively with the abundance of *Anaeralineaceae* (r = −0.854; P ≤ 0.01), *Rhodocycladeae* (r = −0.899; P ≤ 0.01), *Peudomonas* (r = −0.755; P = 0.005) and *Bacteria* (r = −0.804; P = 0.002). The organic matter (OM) correlated positively with the abundance of *Bacillaceae* (r = 0.825; P = 0.001), *Lactobacillus* (r = 0.804; P = 0.002), *Ignatzschineria* (r = 0.771; P = 0.003), *Anaerococcus* (r = 0.76; P = 0.004), *Caldicoprobacter* (r = 0.804; P = 0.002), *Thippseudomonas* (r = 0.902; P ≤ 0.01), *Sinibacillus* (r = 0.872; P ≤ 0.01), *Tepidinicroblum* (r = 0.853; P ≤ 0.01), *Terrisporobacter(r* = 0.755; P = 0.005), and negatively with the abundance of *Anaerolineaceae* (r = −0.858; P ≤ 0.01), *Rhodocycladeae* (r = −0.929; P ≤ 0.01), *Luteimonas* (r = −0.741; P = 0.006) and *Petrimonas* (r = −0.734; P = 0.007). The protease (PE) correlated positively with the abundance of *Clostridiales* (r = 0.825; P = 0.001), *Caldicoprobacter* (r = 0.923; P ≤ 0.01), *Thiopseudomonas* (R = 0.713; P = 0. 009), *Sinibacillus* (r = 0.732; P = 0.007), *Tepidimicrobium* (r = 0.755; P = 0.005), Limnochordaceae (r = 0.767; P = 0.004), *Moheibacter* (r = 0.811; P = 0.001) and *M55-D21(r* = 0.79; P = 0.002), and negatively with the abundance of *Anaerolineaceae* (r = −0.737; P = 0.006). The pH correlated positively with the abundance of the *M55-D21* (r = 0.725; P = 0.008) and *MBA03(r* = 0.76; P = 0.004).

**Fig 6.**
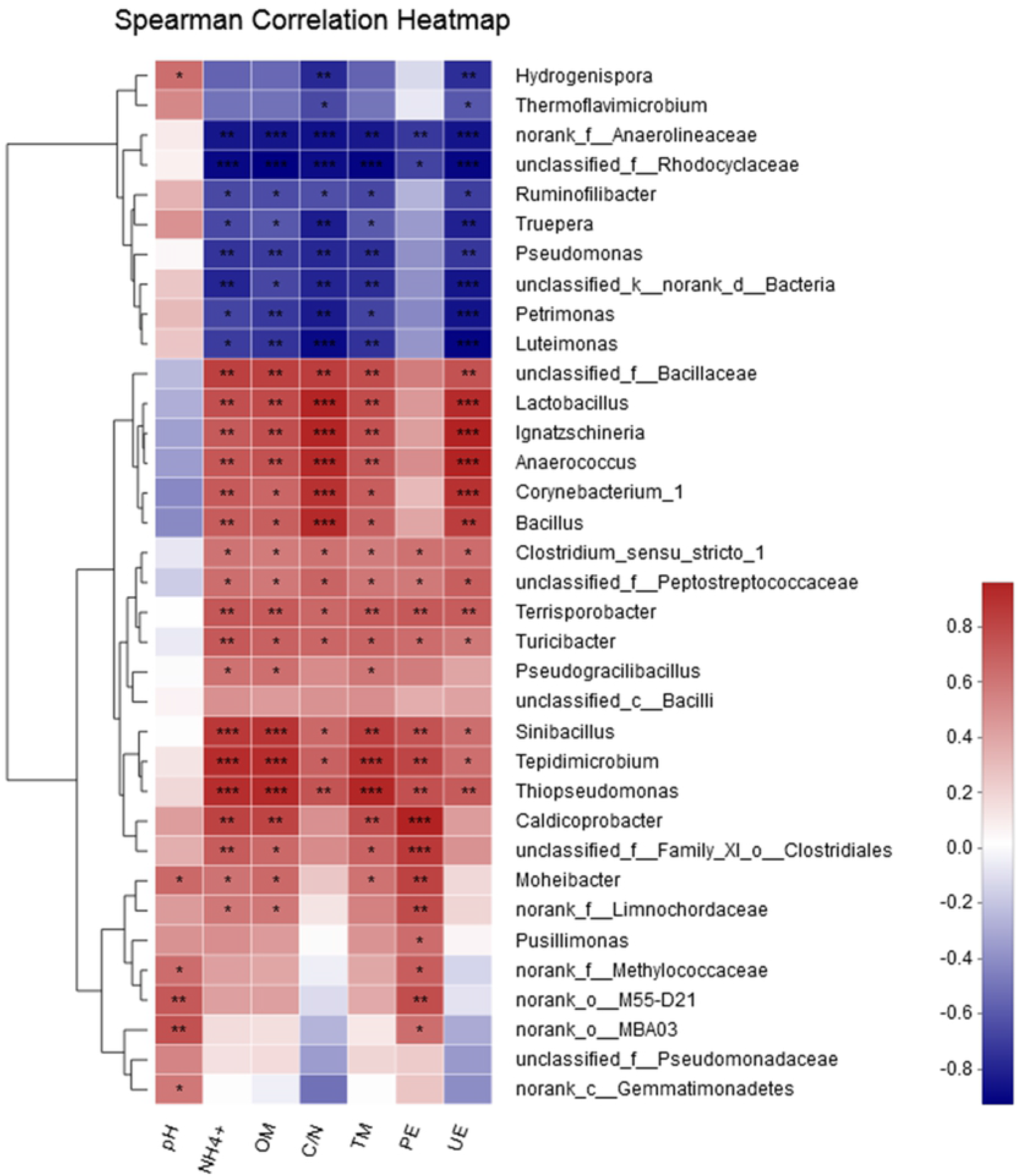
Correlation between Chemical parameters and genus abundance. The top and left hierarchical cluster based on the corresponding correlation matrix between chemical parameters and genus abundance. The similar clusters were found with complete linkage method. Only the predominant bacterial genera (the dominant species of the top 35 in at least one sample) are presented. Cells are colored based on the Spearman correlation coefficient between chemical parameter and genus abundance. The red color represents a positive correlation, and the green color represents a negative correlation. (PE, protease; OM, organic matter; TM, temperature; UE, urease; *, 0.01 < P ≤ 0.05; **, 0.001 < P ≤ 0.01; ***, P ≤ 0.001)

Using traditional methods, Bach and Munch found that *Pseudomonas fluorescens, Bacillus cereus*, *Bacillus mycoides*, *Cytophaga* and *Flavobacterium* wrere present in many soils leading the process of proteins hydrolysis, as they secreted metalloproteinases [39]. Watanabe and Hayano observed that peptide enzymes from *Bacillus*, especially a neutral metal peptidase secreted by *Bacillus cereus* and *Bacillus mycoides*, and also an alkaline serine protease secreted by *Bacillus subtilis*, was the main protease involved in peptide degradation in paddy soil [40]. In this study, the protease in the compost was higher in earlier stage. The bacteria of *Cytophaga* were more abundant in compost samples of maturity stage, whereas *Flavobacterium* was richer in the primary stage and high temperature period of compost, but less in maturity stage of compost. Through correlation analysis, in the early stage of compost the bacteria with important function for improving the compost proteases was mainly from the *Firmicutes* and *Bacteroidetes* phyla. These groups were the main degradation agents of nitrogenous organic compounds in the compost.

In all kinds of bacteria, the secreted proteases were mainly alkaline serine proteases and neutral metalloprotease [41]. The activity of protease could be used as an indicator of ammonification [42]. The change of pH throughout composting was not particularly intense, being maintained between 7.5 and 8.1, and was mainly related to *Firmicutes*.

Organic matter, ammonium nitrogen and temperature had similar influence on bacterial populations, suggesting that these species of bacteria lived mainly in the high-temperature compost period and with a high concentration of ammonium nitrogen environment in the compost. These bacteria growth in the early stage of the compost, when the content of organic matter conditions was rich, thus playing an important role in decomposition of organic matter.

The microbial communities related by carbon to nitrogen ratio and urease levels had similarities. The microbial community of *Corynebacterium, Bacillus, Lactobacillus, Ignatzschineria* and *Anaerococcus* had strong urease activity, and this microbial community intensified the change of carbon and nitrogen ratio in the compost. The nature of compost was the process of microbial degradation of complex organic compounds in a specific environment. Microorganisms change the environment of composting constantly in the process of metabolism. Reciprocally, changes in environmental conditions lead to changes in microbial community structure.

## 3. Conclusions

In this study, the compost was made of mixture of pig manure and wood chips. We found that the protease activity, organic matter content and ammonium nitrogen concentration were higher in the early stage of composting. Meanwhile, the urease activity was highest in the high temperature period. The carbon to nitrogen ratio of the compost decreased continuously with fermentation. The dynamic change in the composition of bacterial overtime in the compost of a 180 kg piles were explored using microbial diversity analysis. The results showed that the microbial species increased with the compost fermentation. At the early stage of composting, *Firmicutes* and *Actinomycetes* were dominant. The microbes in the high temperature period were mainly composed of *Firmicutes* and *Proteobacteria* while the proportion of *Bacteroides* was increased during the cooling period. In the compost of maturity stage, the proportion of *Chloroflexi* increased, becoming dominant species with other microorganisms including *Firmicutes*, *Proteobacteria*, *Bacteroides*, *Chloroflexi* but not *Actinomycetes*. Bacteria involved in lignocellulose degradation, such as those of the *Thermobifida, Cellvibrio, Mycobacterium, Streptomyces* and *Rhodococcus*, were concentrated in the maturity stages of composting. Through correlation analysis, the environmental factors including organic matter, ammonium nitrogen and temperature were consistent with the succession of microbial including *Rhodocyclaceae*, *Anaerolineaceae, Thiopseudomonas, Sinibacillus* and *Tepidimicrobium*. The change of urease activity and carbon to nitrogen ratio corresponded to microbial communities, mainly containing *Anaerolineaceae, Rhodocyclaceae, Luteimoas, Bacillaceae, Corynebacterium, Bacillus, Anaerococcus, Lactobacillus, Ignatzschineria* and *Bacillaceae*.

## Acknowlegements

This work was supported by the Open Research Fund of China Tobacco Yunnan Industrial Co., Ltd., (Grant No. 2017CP01, 2017539200370271), the National Natural Science Foundation of China (Grant No. 21572242), the Talents of High Level Scientifc Research Foundation (Grant No. 6631113326) of Qingdao Agricultural University.

